# A High-quality Oxford Nanopore Assembly of the Hourglass Dolphin (*Lagenorhynchus cruciger*) Genome

**DOI:** 10.1101/2024.05.30.596754

**Authors:** Nick McGrath, Jamie le Roux, Annabel Whibley, Alana Alexander, Ramari Oliphant Stewart, Muriel Johnstone, Karen A. Stockin, Olin K. Silander

## Abstract

The hourglass dolphin (*Lagenorhynchus cruciger*) is a small cetacean species of the Southern Ocean, with significance to iwi Māori (*Māori tribes*) of Aotearoa New Zealand as taonga (*treasured/valued*). Due to the remoteness and difficulty of surveying Antarctic waters, it remains one of the least-studied dolphin species. A recent stranding of an hourglass dolphin represented a rare opportunity to generate a genome assembly as a resource for future study into the conservation and evolutionary biology of this species. In this study, we present a high-quality genome assembly of an hourglass dolphin individual using a single sequencing platform, Oxford Nanopore Technologies, coupled with computationally efficient assembly methods. Our assembly strategy yielded a genome of high contiguity (N50 of 8.07 Mbp) and quality (98.3% BUSCO completeness). Compared to other Delphinoidea reference genomes, this assembly has fewer missing BUSCOs than any except *Orcinus orca*, more single-copy complete BUSCOs than any except *Phocoena sinus*, and 20% fewer duplicated BUSCOs than the average Delphinoidea reference genome. This suggests that it is one of the most complete and accurate marine mammal genomes to date. This study showcases the feasibility of a cost-effective mammalian genome assembly method, allowing for genomic data generation outside the traditional confines of academia and/or resource-rich genome assembly hubs, and facilitating the ability to uphold Indigenous data sovereignty. In the future the genome assembly presented here will allow valuable insights into the past population size changes, adaptation, vulnerability to future climate change of the hourglass dolphin and related species.

## Introduction

The hourglass dolphin (*Lagenorhynchus cruciger)* is a small cetacean species that inhabits pelagic Antarctic and sub-Antarctic waters. Although its northernmost range includes Aotearoa New Zealand, where it is considered a taonga (*treasured)* species by iwi Māori (*Māori tribes*), the hourglass dolphin is strongly associated with the Antarctic Convergence, and rarely found in close proximity to land masses (Acevedo, Garthe, and González 2017; Dellabianca et al. 2012; Santora 2012). Although not an uncommon species (Goodall et al. 1997), due to the remoteness and difficulty of surveying Antarctic waters, it is one of the least studied species of dolphin (Dellabianca et al. 2012; Goodall et al. 1997), with information largely limited to observations from living animals: group size, locality, acoustics, and presence of calves (Goodall 1997; Acevedo, Garthe, and González 2017; Dellabianca et al. 2012; Tougaard and Kyhn 2009; Kyhn et al. 2009; Thiele, Chester, and Gill 2000; Todd and Williamson 2022).

For an enigmatic oceanic species such as the hourglass, the few beach stranding events that have occurred have provided important opportunities to learn more about the physiology and other phenotypes such as diet preference (e.g. (Goodall et al. 1997; Fernandez et al. 2003; Marchesi et al. 2016; Peters et al. 2022). Physical specimens from a stranding also provide an opportunity to obtain genetic material for genome sequencing. These genomes can be a pathway towards understanding past population size changes, adaptation, and vulnerability to changing climate - an aspect particularly important for the hourglass dolphin in the rapidly changing Antarctic environment (MacLeod 2009). In addition, considerable uncertainty remains around the evolutionary relationships between Lissodelphininae - the subfamily of true dolphins to which the hourglass dolphin belongs (Goodall et al. 1997; Vollmer et al. 2019; Harlin-Cognato and Honeycutt 2006; Cope 1866; McGowen 2011; Banguera-Hinestroza et al. 2014; Leduc, Perrin, and Dizon 1999). Genomic data could be particularly useful in resolving such taxonomic confusion, as demonstrated recently in resolving the taxonomic placement of the pygmy right whale (*Caperea marginata*) (Dutoit et al. 2022).

However, the generation and analysis of genomic data can exacerbate pre-existing inequities around who has access to the technology to generate such genetic resources, who gets to decide how these resources are looked after, and who benefits from the generation of such resources (Mc Cartney et al. 2022; Te Aika et al. 2023). This situation is particularly heightened when it comes to sequencing species that are taonga (*treasured*) by Indigenous peoples. In these cases, the tikanga (*protocols*) around protecting and ensuring the safety of tissues and genetic resources is paramount for upholding Indigenous data sovereignty (Robbins et al. 2023; Jennings et al. 2023).

One pathway toward upholding Indigenous data sovereignty, and ensuring that capacity-building can occur within the communities where rare species are found, is to develop methods for sequencing and genome assembly that are achievable locally. Here, we provide evidence that this is currently possible for mammalian-sized genomes. We present a genomic assembly for an individual hourglass dolphin that relies solely on sequence from the Oxford Nanopore Technologies sequencing platform. We compare multiple basecallers (Guppy and Dorado) and assemblers (Raven, Nextdenovo, and Goldrush), obtaining the most contiguous genome using the Dorado basecaller and the computationally efficient Raven assembler. For most mammalian species, obtaining data such as that utilised in our study is achievable on a single Oxford Nanopore PromethION flow cell for approximately $1500 USD. With the use of a low-overhead RAM assembler such as Raven it is possible to implement the complete assembly pipeline on a high-end gaming laptop (although close to 128 Gb of RAM is required and this is available only on high-end laptop computers). Furthermore, we show that this assembly is, on average, more complete than any other published Delphinoidea genome (fewer missing BUSCOs and more complete single copy BUSCOs), including the highly curated bottlenose dolphin (*Tursiops truncatus*).

While reduced capital outlay and sequencing costs have already allowed for a more equitable distribution of the capability to generate genomic resources, this has rarely extended to include organisms with large or complex genomes. The results here suggest that current genomic technologies can be applied to enhance the ability of Indigenous communities to maintain sovereignty over tissues and data collected from their taonga (*treasured*) species, including rarely sampled species such as the hourglass dolphin.

## Methods and Materials

### Specimen Collection

On the 5th of August 2020, an individual hourglass dolphin was reported beached at Orepuki Beach in Te Waewae Bay, Murihiku, Aotearoa New Zealand. This individual was given the customary name Hārua-tai, reflecting its connection to the rough seas of the Southern Ocean. With the permission and full blessing of the tribal authority, Ōraka-Aparima Runaka, we retrieved and necropsied the dolphin on the 29th of September to recover a full suite of samples from major organs including the heart. The customary name, Hārua-tai, was used in addition to a scientific coding system in order to link the whakapapa (*connections*) of the materials recovered from this animal. Cardiac tissue was stored at −80°C until permission from Ōraka-Aparima was granted to undertake this co-designed study to assemble the genome of the hourglass dolphin.

### DNA Isolation and Sequencing

We divided the cardiac tissue into two subsamples (21 mg and 25 mg) and isolated DNA from each using the Monarch Genomic DNA Purification Kit. We lysed the tissue for a total of 6.5 hours, agitating at 600 rpm for the first 30 minutes and 300 rpm thereafter. During both these lysis-agitation steps, we incubated the sample at 56°C. We prepped each DNA sample for Oxford Nanopore sequencing using the ligation sequencing kit (SQK-LSK114) according to the manufacturer’s instructions. We sequenced the samples on two sequential days on two PromethION R10.4.1 flow cells on a P2 Solo instrument for 68.5 hours and 72 hours. We stored the DNA at −20C for three months and performed a third sequencing run using the latter sample, which had exhibited a higher read N50 (7.2 kilobase pairs (kbp) vs. 9.5 kbp).

### Bioinformatics Pipeline Overview

We implemented a straightforward strategy for basecalling and filtering, assembly, assessment of assembly contiguity and completeness, haplotig purging polishing, annotation, and variant calling and phasing (**Fig. 1**).

**Figure 1.**
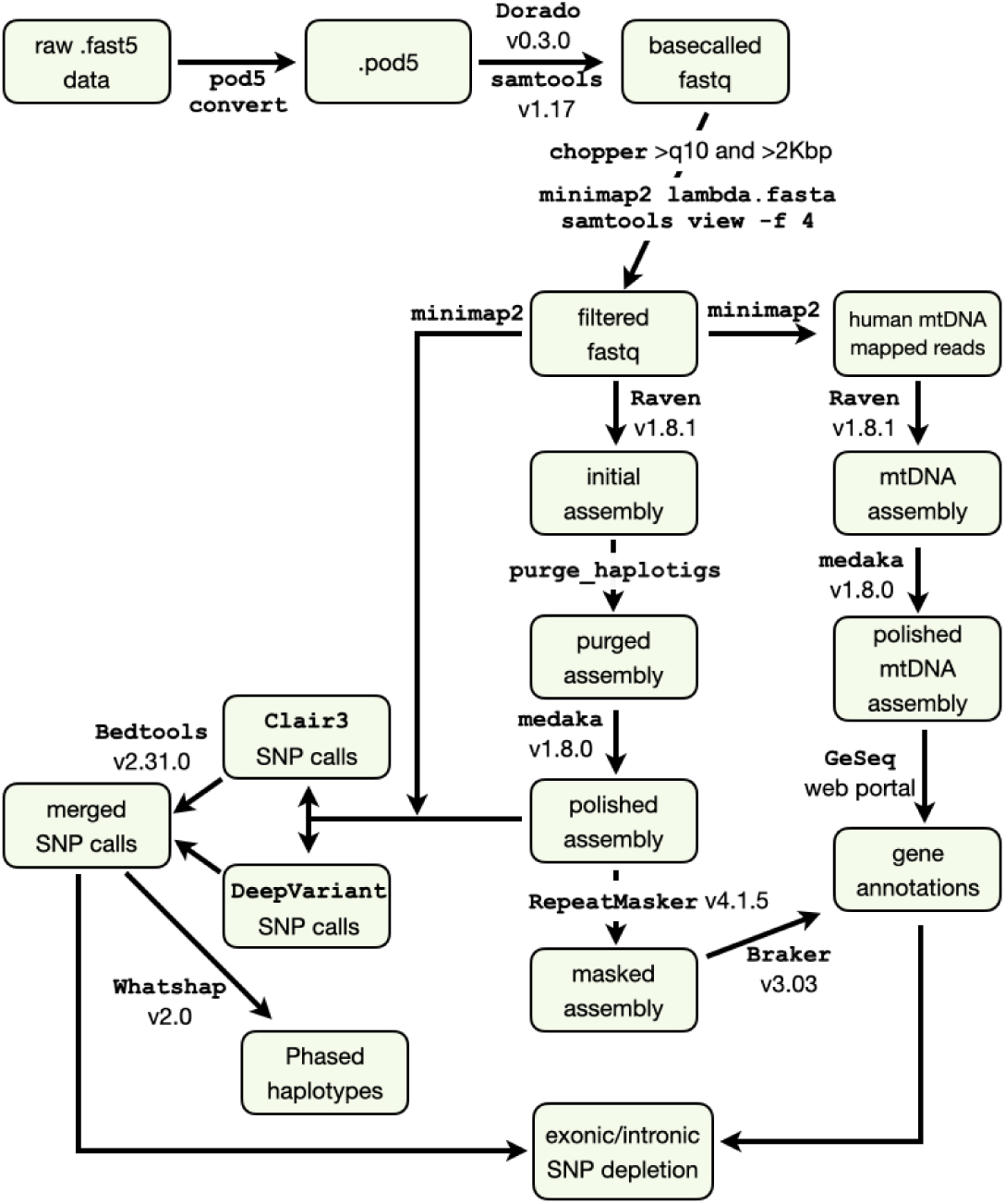
Flowchart of Raven-based assembly. Each software program is noted in Courier serif font; boxes show the results of each step; see methods for details on versions and specific arguments.

### Basecalling and Filtering

We basecalled the fast5 files using *Guppy* v.6.3.9. with the dna_r10.4.1_e8.2_400bps_sup model. We also converted fast5 files to pod5 files using *pod5 convert* from the pod5 package of tools. We basecalled the pod5 files using *dorado* v0.3.0 (*Dorado* 2023) and the dna_r10.4.1_e8.2_400bps_sup@v4.1.0 model and converted the resultant .bam file to .fastq format using *samtools fastq* v1.17 (Danecek et al. 2021). To remove control DNA, we first mapped all reads to the control lambda phage genome using *minimap2* (Li 2018) and removed all mapped reads using *samtools view -f 4*. We filtered the fastq files using *chopper* (De Coster and Rademakers 2023) to retain only reads with average quality scores above 10 and lengths greater than 2 kbp. We also cropped 50 bp from the head and tail of each read. Using 16 threads, this process took 20 minutes and minimal RAM. For downsampling the read data to 90Gb, we used filtlong (Wick and Menzel 2017), prioritising read length (--length_weight 10).

### Assembly

We then assembled both the Dorado-called and Guppy-called data using three different assemblers with RAM requirements less than 400 GB, and which are optimised for Oxford Nanopore data: Goldrush v1.0.1 (Wong et al. 2023), Nextdenovo v2.5.2 (Hu et al. 2023), and Raven v1.8.1 (Vaser and Šikić 2021). Using Raven with the Dorado data took a total time of 16 hours and a maximum of 153 Gb RAM, averaging approximately 130 Gb of RAM (**Supp. Fig. 1**)

For mitochondrial genome assembly, we mapped all reads to the human mtDNA sequence, filtered those reads using chopper (De Coster and Rademakers 2023) to include only reads between 14Kbp and 17Kbp (the approximate size of the full length mitochondrial genome), and subsampled this filtered set using *SeqKit* (Shen et al. 2016) to include only 5% of the reads (due to the extremely high coverage). We assembled these reads using Raven.

We calculated NG50 using Quast v5.2.0 (Gurevich et al. 2013), assuming a genome size of 2.384 Gbp.

### Haplotig Purging and Assembly Polishing

We used purge_haplotigs (Roach, Schmidt, and Borneman 2018) to remove contigs from the assembly that had lower than expected coverage or which were highly similar to other contigs; these were judged as being haplotigs.

We used medaka v1.8.0 (*Medaka* 2023) to polish the assembly using the *r1041_e82_400bps_sup_v4.1.0* model.

In the most contiguous assembly, Dorado-basecalled data assembled with Raven, we found a short contig of 1,396 bp, with the next smallest contig at almost 25 Kbp). Upon blasting this sequence we found it was 99% identical to a sequence from *Bos taurus*. We removed this contig from the assembly, a likely contaminant from other lab samples. We also used Kraken2 (Wood et al. 2019) to determine the taxon assignment for all other contigs in the assembly and found that all were vertebrate in origin, suggesting no microbial contamination. In addition, given the above contaminant *B. taurus* contig, we used mash (Ondov et al. 2016) to calculate approximate distances for each contig to the reference *Bos taurus* genome and the *Tursiops truncatus* genome using 500,000 min-hashes. All contigs were closer to *T. truncatus* than *B. taurus*, although 84 contigs were too small to accurately measure distance using mash. In these cases we mapped the contigs to each reference genome using minimap2 -x lr:hq (Li 2018). For each contig, we found the longest aligned region to either genome and tested whether the *B. taurus* or the *T. truncatus* alignment was longer. The *T. truncatus* alignments were longer except for three contigs with short aligned regions. To test the identity of these contigs, we blasted them against the nr database. The three contigs ranged from 93% to 97% identity to other Delphinid genomes. Thus, we concluded that no other contigs besides the short 1,396 bp above were contaminants.

To map the correspondence of the *L. cruciger* hourglass assembly contigs to the autosome and sex chromosomes of the *T. truncatus* assembly, we aligned the *L. cruciger* assembly to the *T. truncatus* assembly using quarTeT Assembly Mapper (Lin et al. 2023).

### Repeat Regions

We masked repeats using RepeatMasker v4.1.5 (Bogdahn 2015) with the Dfam v3.7 repeat element database, nhmmscan version 3.3.2, and taxa search limited to mammals. We identified telomeric repeats (TTAGGG) using SeqKit (Shen et al. 2016). To calculate kmer repetition in contigs we used jellyfish 2.2.10 (Marçais and Kingsford 2011). For each contig, we counted the frequency of all 21-mers, and then calculated the fraction of 21-mers that were present once vs. more than once. We used SeqKit (Shen et al. 2016) to count N content across regions masked by RepeatMasker.

### Assembly Completeness

We assessed assembly completeness using compleasm.py (Huang and Li 2023) and the odb10 versions of five databases: Laurasiatheria, Cetartiodactyla, Eutheria, Mammalia, and Eukarya. We compared the assembly here to nine additional Delphinoidea assemblies (**Supp. Table 1**) to which we applied compleasm in the same way.

### Annotation

We annotated the masked genome using Braker v.3.0.3 (Hoff et al. 2016; Brůna et al. 2021; Hoff et al. 2019; Brůna, Lomsadze, and Borodovsky 2020; Buchfink, Xie, and Huson 2015; Gotoh 2008; Lomsadze et al. 2005; Stanke et al. 2008; Iwata and Gotoh 2012; Stanke et al. 2006) and the OrthoDB11 Vertebrate database of protein families (March 3 2023) (Kuznetsov et al. 2023) downloaded from https://bioinf.uni-greifswald.de/bioinf/partitioned_odb11/.

We annotated the mitochondrial genome using GeSeq (Tillich et al. 2017) with the BLAT reference sequences set to all Delphinid mtDNA genomes in RefSeq.

### Variant Calling and Phasing

We used Clair3 v1.0.4 (Zheng et al. 2022) to call variants using the Clair3 *r1041_e82_400bps_sup_v410* model. We observed a clear dip in the quality of SNV calls at a quality score of 14 (**Supp. Fig. 2**), and using *rtg vcffilter* v3.12.1 (Cleary et al. 2014), we filtered the calls to include only those with qualities above 14. We also used DeepVariant (Poplin et al. 2018) with *--model_type=ONT_R104*. In this case there was a clear dip in the quality of calls at a quality score of 20 (**Supp. Fig. 2**), and retained only calls with scores above that. Finally, we obtained a set of high quality calls by intersecting these two call sets using *bcftools* v1.17 (Danecek et al. 2021) *norm* and *isec*. We then used bedtools v2.31.0 (Quinlan and Hall 2010) to find intersections between individual bed files of genic, exonic, and intronic regions and these variant calls.

We phased variant calls using *Whatshap* v2.0 (Martin et al. 2016). To visualise the phasing we used *whatshap haplotag* and IGV v2.1.62 (Thorvaldsdóttir, Robinson, and Mesirov 2013).

## Results

Using Dorado, overall we obtained 8.51 million reads and 39.6 Gbp; 4.96 million reads and 31.3 Gbp; and 20.4 million reads and 110.2 Gbp of data from the three Oxford Nanopore runs, respectively. The N50 values for these runs were 7.2 kilobase pairs (Kbp), 9.5 kbp, and 8.4 kbp. After pooling these three runs and filtering, using Dorado, 20.3 million reads and 142.6 Gbp remained, with an N50 of 9.04 kbp; using Guppy and filtering the pooled data resulted in 21.1 million reads and 145.9 Gbp, with an N50 of 8.88 kbp. Using both datasets and all three assemblers (Goldrush, Raven, and Nextdenovo, we found that contiguity differed considerably (**Table 1**), with the Dorado-basecalled Raven assembly having the longest contig NG50 and maximum contig length (8.08 Mbp and 39.1 Mbp, respectively. Both of these are almost 50% longer than the least contiguous method, Guppy with Goldrush.

**Table 1.** Assembly contiguity across basecallers and assemblers. The statistics below are for assemblies before polishing, haplotig purging, or contaminant removal.

| Assembler | Basecaller | Number of contigs | Total length (Gbp) | Min. contig length | Max. contig length (Mbp) | NG50 (Mbp) |
| --- | --- | --- | --- | --- | --- | --- |
| Goldrush | Guppy | 40,014 | 2.57 | 1 | 28.7 | 5.68 |
| Goldrush | Dorado | 29,863 | 2.44 | 1 | 24.5 | 6.29 |
| Nextdenovo | Guppy | 1,090 | 2.32 | 15,220 | 28.4 | 6.55 |
| Nextdenovo | Dorado | 1,098 | 2.32 | 16,390 | 27.5 | 6.42 |
| Raven | Guppy | 1,289 | 2.45 | 1,380 | 24.9 | 7.32 |
| Raven | Dorado | 1,243 | 2.44 | 1,379 | 39.1 | 8.08 |
| Raven | Dorado Q16 | 1,187 | 2.40 | 10,313 | 30.6 | 6.38 |

We obtained 31 Mbp of reads (subsampled to include only 5% of total number) between 14 Kbp and 17 Kbp that mapped to the human mitogenome, with an average length of 15,923 bp. Using Dorado-basecalled data and Raven (the most successful whole-genome combination), we assembled a circular 16,389 bp mitochondrial contig. This was 96.9% identical to the 16,392 bp Pacific white-sided dolphin *L. obliquidens* mitogenome (Lee et al. 2018) and 96.6% identical to the 16,371 bp Heaviside’s dolphin *C. heavisidii* mitogenome (Hassanin et al. 2012), the two closest matches in the NCBI database.

### Assembly Completeness Across Assemblers

We quantified assembly completeness using compleasm (Huang and Li 2023), a new implementation of BUSCO (Simão et al. 2015) using the Laurasiatheria odb10 database. Again we found that the Dorado-basecalled Raven assembly had the highest number of complete single copy BUSCOs and the fewest fragmented or missing BUSCOs compared to the other five baseball-assembly combinations. (**Supp. Table 2**). Due to the Dorado-based Raven assembly having the highest contiguity and BUSCO completeness, we used this assembly as the basis for the remainder of the analyses here.

### Haplotig Purging and Assembly Polishing

We performed a single round of polishing and purging of haplotigs, resulting in a final assembly 2.384 Gbp in length, with 894 contigs, an NG50 of 8.074 Mbp, an L50 of 89, a maximum contig length of 39.03 Mbp, and a minimum contig length of 24,985 bp. This assembly size is similar to the total chromosomally-scaffolded portions of other delphinid species such as the bottlenose dolphin *Tursiops truncatus* (2.343 Gbp), the common dolphin *Delphis delphinus* (2.364 Gbp), the white-beaked dolphin *Lagenorhynchus albirostris* (2.404 Gbp) and the long-finned pilot whale *Globicephala melas* (2.364 Gbp). However, it is considerably smaller when including the unplaced scaffolds of these assemblies (2.637, 2.774, 2.767, and 2.651 Gbp, respectively). This may be due to the small repetitive contigs in this *L. cruciger* assembly being artifactually collapsed.

To estimate the copy number of each contig, we mapped the raw reads back onto the assembly and filtered the mapped reads to include only those with a mapping quality greater than 20. We found that non-repetitive contigs (see below) had largely bimodal read depths, with the majority having a depth of approximately 52 and a smaller number having a depth of approximately 26 (**Fig. 2**), which we inferred were contigs belonging to the X and Y chromosomes in this male individual. We confirmed this by aligning the contigs to the *T. truncatus* genome assembly (a female individual). We found that out of the 129 non-repetitive contigs with median depth less than 30, 112 aligned to the X chromosome, four did not align, ten aligned to unplaced scaffolds, and three aligned to autosomes.

**Figure 2.**
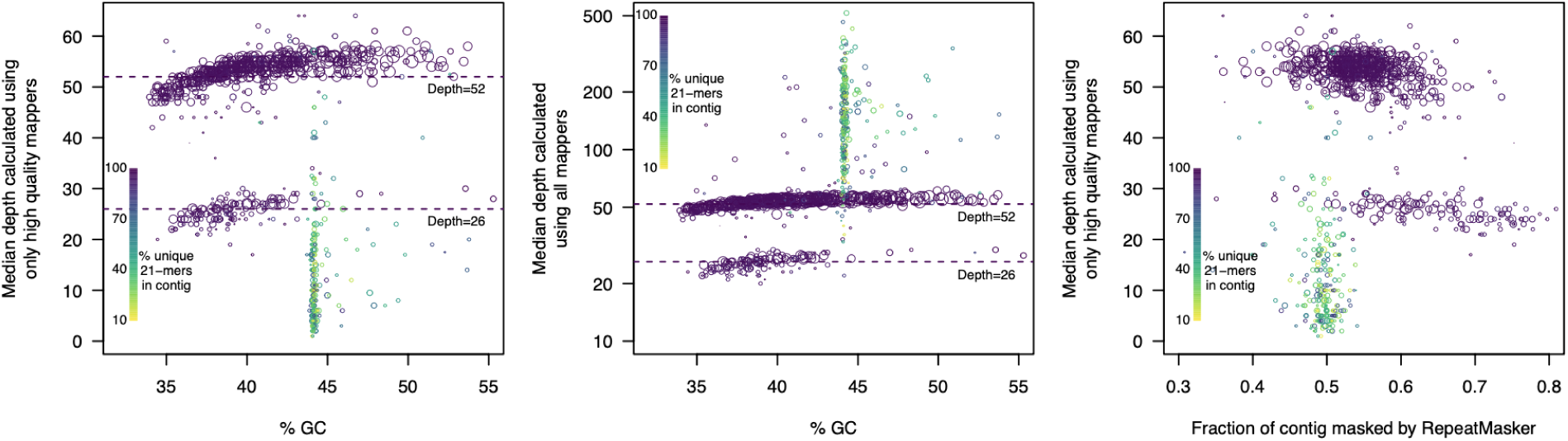
Sequencing depth, GC content, and repetitiveness varies across contigs. A. GC-content vs. depth calculated using high quality mappers. Each contig is represented by a circle that is scaled to the log of contig size. Depth was largely bimodal, with the lower mode (approximately 26) corresponding to the X and Y chromosomes in this male individual. A slight bias toward lower depth in very GC-poor contigs was apparent (visible as an upward slope as % GC increases). A small number of contigs exhibited very similar %GC values (approximately 44%); these contigs had low nucleotide diversity as measured by kmer diversity (green and yellow circles). The low nucleotide diversity of these contigs also resulted in very few high quality mappers and thus low depth. **B**. **GC-content vs. depth calculated using all mapped reads.** The same plot as in (**A**) but when all mapped reads are included. In this case, contigs with low nucleotide diversity exhibited extremely high depth, suggesting that most arose from hard-to-resolve genomic regions that were collapsed during assembly. Note that the scale of the y-axis is linear for (**A**) and logarithmic for (**B**) due to the high coverage values for some repetitive contigs (up to 500-fold). **C. Masked contig fraction vs. depth**. Colours are identical to (**A**) and (**B**). The x-axis indicates the fraction of each contig that was designated by RepeatMasker as containing repetitive elements (e.g. LINEs, SINEs, LTRs) and thus masked with Ns. Contigs that have low nucleotide diversity are depleted for repetitive genomic elements (green points), with most having approximately 50% repetitive element content as opposed to almost 60% for most autosomal contigs.

However, the *T. truncatus* is a female individual and has no Y chromosome. Of the four unaligned contigs (176 kbp, 103 kbp, 74 kbp, and 70 kbp in length), the top blast hits by bit score were, respectively: *Bos taurus* chr Y (80% identity across the alignment, but 93% identity to *D. delphis* chr 12); *G. melas* chr 19 (95% identity); *Balaenoptera acutorostrata* chr 19 (81% identity), and *D. delphis* chr Y (92% identity). Thus, it appears that some unmapped contigs in this hourglass assembly are homologous to other species’ Y chromosomes. The match of a single contig with both the *B. taurus* Y chromosome and *D. delphis* chr 12 suggests that other assemblies may have sex chromosome contigs assigned to autosomal contigs.

A number of contigs had lower coverage of high-quality mapped reads, as well as GC percentages in a very narrow range (**Fig. 2**). These contigs also tended to be smaller (less than 500 kbp). When we quantified nucleotide diversity in these using kmer content, we found that it was extremely low (**Fig. 2A**), with the majority having less than 50% of all 21-mers being unique (**Methods**). For this reason, we suspected that they had been artificially collapsed during assembly, resulting in very few reads mapping with high quality. As expected, when we calculated depth based on all mapped reads rather than just high quality mappers, these contigs exhibited very high coverage (**Fig. 2B**). Upon alignment to the *Tursiops truncatus* genome, which is one of the most well-curated delphinoid assemblies, we found that 181 out of the 254 highly repetitive contigs (unique kmer content less than 90%) mapped to unplaced *T. tursiops* scaffolds, and were enriched for the telomeric motif TTAGGG (Zhong et al. 1992): 235 out of 254 were more than four-fold enriched for this motif compared to only 16 of the 640 non-repetitive contigs. This suggests that some of these are artifactually collapsed subtelomeric or telomeric regions.

To further characterise these problematic contigs, we examined the repeat element content. We first used RepeatMasker to determine repetitive genomic elements. 51% of the genome was designated as consisting of repetitive elements, including 111 Mbp of SINE elements (4.68% of the total genome), 704 Mbp of LINE elements (29.6%), and 236 Mbp of LTR elements (9.90%). Notably, the low nucleotide diversity contigs (few unique kmers) were depleted for repetitive elements (**Fig. 2C**). This supports the hypothesis that these are not collapsed due to repetitive elements; rather they are collapsed due to extremely low nucleotide diversity and are possibly telomeric, centromeric, or parts of the Y chromosome.

### Annotation

We annotated the repeat-masked genome with Braker3, resulting in 129,464 exons and 104,187 introns with mean lengths of 157 bp and 1,509 bp, respectively. 25,329 protein coding genes in total were annotated, with an average length of 268 amino acids and a maximum length of 4,065 amino acids. This number of protein coding genes is considerably larger than the number annotated in *T. truncatus* (19,240) despite this assembly having lower BUSCO scores. This is most likely due to differences in annotation methods, as *T truncatus* was annotated using the NCBI eukaryotic genome annotation pipeline, which relies on Prosplign and Gnomon.

The annotated mitochondrial genome had the expected 22 tRNAs, 12S and 16S ribosomal RNAs, and 13 coding sequences.

### Completeness of Final Assembly

To check the accuracy and completeness of the final polished and purged assembly, we repeated the compleasm analysis. The final assembly exhibited slightly improved BUSCO scores in Laurasiatheria (98.37%) and averaged 98.3% single copy complete BUSCOs across five different lineages (**Supp. Table 3**).

We compared the completeness of the hourglass assembly to nine other Delphinidae assemblies, which we selected on the basis of their being the most contiguous and complete assemblies in this family (all are designated as reference genomes by NCBI). However, we also excluded several genomes designated as reference quality as they had far fewer single copy complete BUSCOs, including *Steno bredanensis*, *Grampus griseus*, and *Sousa chinensis*. We found that the hourglass assembly had more complete and single copy BUSCOs (12,056; **Table 2**) than any other assembly except the vaquita, *Phocoena sinus* (12,072); correspondingly it had fewer missing BUSCOs (48) than any other assembly except the orca, *Orcinus orca* (41). 33 of these BUSCOs were missing from all Delphinoidea assemblies (**Fig. 3**), and were likely lost in the ancestral lineage. If these were indeed lost, this suggests the quality of the hourglass assembly is even higher than first appears: excluding the 33 BUSCOs missing in all, while the hourglass is missing only 15 out of 12,201, the average reference-level Delphinoidea assembly is missing 26.

**Figure 3.**
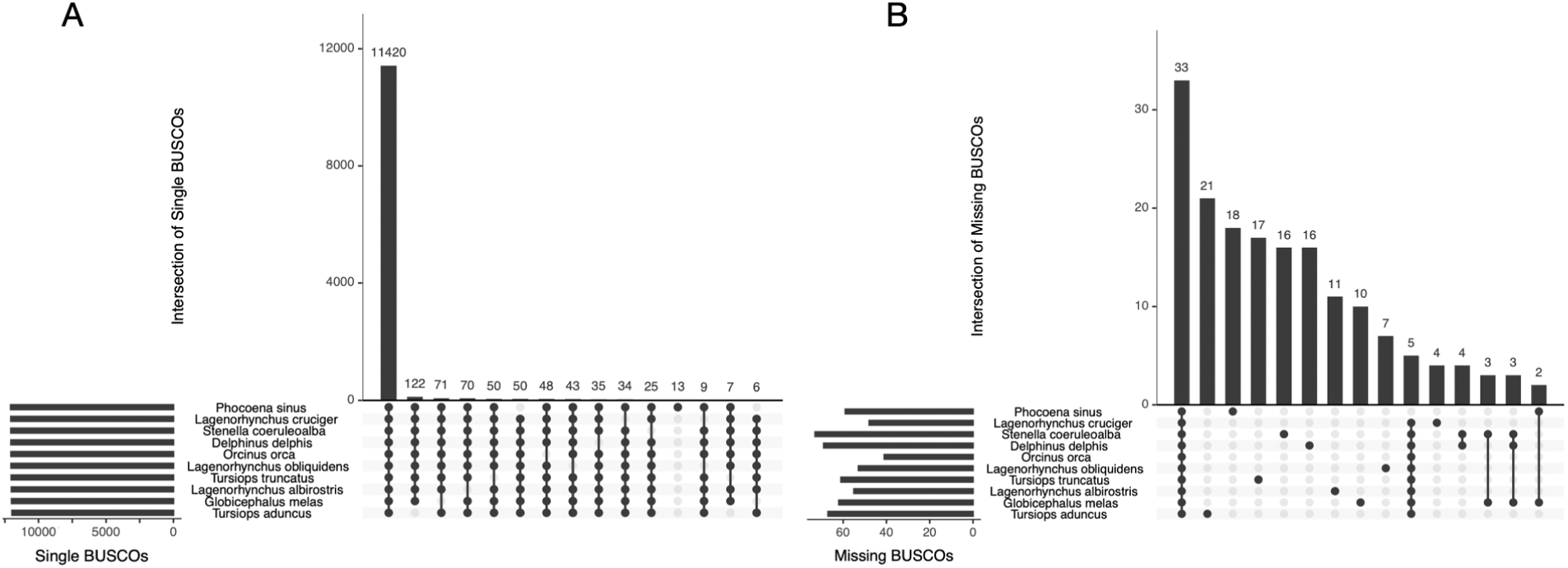
Upset plots of BUSCO overlaps across Delphinoidea assemblies. A. Intersection of single copy and complete BUSCOs using the Laurasiatheria ODB10 database. Each black point on a vertical line indicates taxa that share a set of BUSCOs, while each grey point indicates that the set of BUSCOs is missing from that taxon. The top bars show the total number of BUSCOs in each set (for example, 11,420 BUSCOs are present as single copy-complete in all ten taxa; 122 are single copy-complete in all taxa except *T. aduncus*). The left bars indicate the total number of single copy complete BUSCOs in each taxon. **B. Intersection of missing BUSCOs.** As in (A), overlaps in BUSCO sets are indicated by black points and taxa that are not part of that set are in grey. 33 BUSCOs are missing from all Delphinoidea, most of which were likely lost in the ancestor of these taxa. For both plots, only the 15 intersections with the most BUSCOs are shown, as there are a large number of intersections that contain only one or two BUSCOs.

**Table 2.** Assembly completeness across Delphinoidea. Each column indicates the total number of single, duplicated, missing, or fragmented BUSCOs as found by Compleasm.

| Species | Single copy | Duplicated | Missing | Fragmented |
| --- | --- | --- | --- | --- |
| *Phocoena sinus* | 12072 | 80 | 59 | 23 |
| *Lagenorhynchus cruciger* | 12056 | 96 | 48 | 34 |
| *Stenella coeruleoalba* | 12055 | 87 | 73 | 19 |
| *Delphinus delphis* | 12049 | 94 | 69 | 22 |
| *Orcinus orca* | 12048 | 129 | 41 | 16 |
| *Lagenorhynchus obliquidens* | 12043 | 112 | 53 | 26 |
| *Tursiops truncatus* | 12028 | 119 | 61 | 26 |
| *Lagenorhynchus albirostris* | 12013 | 135 | 55 | 31 |
| *Globicephala melas* | 12002 | 148 | 62 | 22 |
| *Tursiops aduncus* | 11969 | 157 | 67 | 41 |

The number of duplicated BUSCOs in the hourglass was 20% lower than average (96 vs. 116); however, the number of fragmented BUSCOs was 30% higher than average (34 vs. 26). This may be an indication that the Oxford Nanopore data results in an increased number of uncorrected small insertions and deletions resulting in truncated reading frames, compared to the combination of PacBio and Illumina data used for most of the other assemblies (**Supp. Table 1**).

We found that the BUSCO results correlated with patterns expected from the evolutionary relationships of the taxa. 50 BUSCOs were present as single copy and complete in all taxa except the most diverged taxon, the vaquita; similarly, the vaquita contained 13 single copy and complete BUSCOs that were not single copy and complete in all other taxa (**Fig. 3**). Nine single copy complete BUSCOs were not single copy-complete in both the hourglass and *Lagenorhynchus obliquidens*, the Pacific white-sided dolphin, but were present as single copy-complete in all others. The hourglass and Pacific white-sided dolphin are likely sister species to the exclusion of *Lagenorhynchus albirostris*, although additional phylogenetic analyses are needed to confirm this. Similar patterns were apparent for both *Tursiops* sister species.

### Variant Calling and Phasing

We called heterozygous sites using both Clair3 and DeepVariant. These have been shown to be the most accurate Oxford Nanopore variant callers, with median indel F1 scores above 99.5% and SNP F1 scores at or above 99.99% at 50X depth in bacterial genomes (Hall et al. 2024).

Clair3 called 4,848,349 million SNVs, 620,899 insertions, and 516,291 deletions. When we filtered these to qualities above 14, this resulted in 4,223,575 SNVs, 208,561 insertions and 278,677 deletions. DeepVariant called 6,869,645 SNVs, 1,069,178 insertions and 3,381,036 deletions, which when filtered to those with quality 20 or more, resulted in 4,188,921 SNVs, 327,370 insertions, and 287,574 deletions. Overlapping the two call sets resulted in a total of 4,021,582 high quality SNPs, 181,409 insertions, and 246,147 deletions (note that the designation of calls as insertions or deletions arbitrarily depends on the state of these positions in the haploid genome assembly).

We also examined the distribution of polymorphic sites across the genome. We hypothesised that due to selection, exons should harbour fewer SNPs relative to introns and intergenic regions, and that exons should harbour few indels. Of the high confidence SNP calls, 0.551% were within exons. This contrasts with the 0.853% of the genome that is annotated as exonic. 6.83% SNPs calls were within introns, closely matching the 6.60% of the genome that is intronic. For indels, only 0.245% lay within exons, a 3.5-fold depletion compared to genome-wide. In addition, exons were depleted for indels that were not multiples of three bp in length; this was not true for introns (**Supp. Fig. 3**). Surprisingly, we observed little depletion for 1 bp indels in exons. These may be due to miscalls present in Oxford Nanopore data at homopolymeric runs, or to segregating slightly deleterious mutations. Additional analyses of the locations, types, and functional effects of these indels would be required to yield insight into this.

Finally, we used Whatshap (Shafin et al. 2021) to phase the genome. 55% of all contigs were phased into a single block, and 80% were phased into three or fewer blocks (Fig. 4). Thus, despite relatively low levels of polymorphisms (0.1%) in many contigs (Fig. 4C and 4D; **Supp. Fig. 4**), successful phasing was possible. However, without additional information, it is difficult to say whether there was a substantial amount of haplotype switching in these phased blocks.

**Figure 4.**
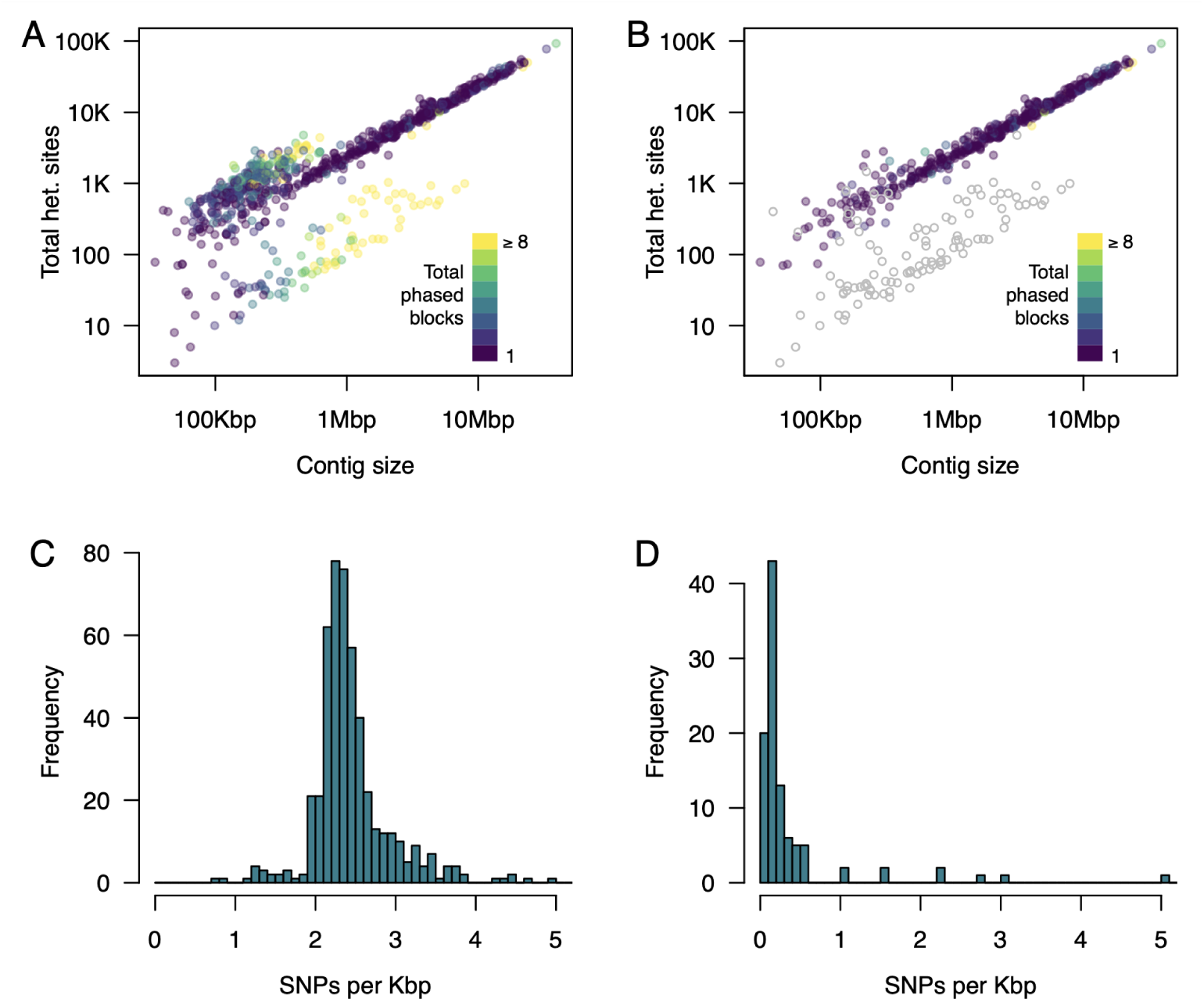
The majority of contigs are phased into a single block. **A**. Number of phase blocks per contig relative to contig size and the number of heterozygous sites in each contig. The majority of contigs that are not fully phased into a single block have low levels of polymorphism (green and yellow points) except for a small number of very large contigs that could not be fully phased (upper left points above the diagonal). **B**. The same plot as in panel (A) but with repetitive contigs removed (defined here as contigs with less than 90% unique 21-mers and putative sex chromosome contigs (coverage less than 30) coloured in grey. The sex chromosome contigs are inferred as having approximately 20-fold lower levels of polymorphism, likely resulting from a combination of assembly errors, false positive calls, and the homologous pseudoautosomal regions in the X and Y chromosomes. **C**. The number of SNPs per Kbp in non-repetitive contigs (unique 21-mers greater than 90%) and with greater than 45-fold coverage of high quality mappers (presumed autosomal contigs). Most autosomal contigs have approximately 2.4 heterozygous SNPs per Kbp with very little deviation from this average. D. The number of SNPs in non-repetitive contigs with less than 30-fold depth (presumed sex chromosome contigs). These contigs averaged 0.11 SNPs per Kbp.

## Conclusions

The genome assembly we present here is one of the most complete of any cetacean, with BUSCO completeness above 98%. Despite relying solely only on Oxford Nanopore data, this quality exceeds many recent cetacean assemblies, even those relying on multiple sequence technologies, most frequently PacBio and Illumina (**Table 2**; **Supp. Table 1**).The use of the Raven genome assembler decreased the required computational resources to just above the size of the read dataset (close to 150 Gb RAM for the 142 Gbp dataset). The required RAM is easily decreased by decreasing the size of the data set (e.g. 98 Gb RAM for a 90 Gbp dataset (**Supp. Fig. 1**). This decreases both sequencing and compute costs while not severely compromising assembly quality, although a full assessment of the effects of changes in coverage and filtering is beyond the scope of what we present here. Low compute cost with few memory limitations is also available via the cloud. However, for the particular assembly presented here, the feasibility and desirability for this as a solution is dependent on how indigenous data sovereignty is implemented: for example, here and in other cases there may be a strong desire to ensure all data are collected and analysed within country or even more locally.

In 2020, the Vertebrate Genome Project estimated $20,000 USD for a mammalian genome, a two-fold reduction from an estimated $40,000 USD in 2018 (Morin et al. 2020). Here we present a high quality genome, from DNA to curated assembly, for less than a tenth of that price, and consuming considerably less computational resources. Overall, the results here illustrate the potential for inexpensive, and more importantly, accessible, assembly of large genomes, enabling community participation and data sovereignty (Garrison et al. 2019).

Finally, this high-quality genome should assist considerably in resolving taxonomic uncertainties in the subfamily Delphininae. In addition, by providing a comprehensive genomic resource, this study will contribute to a deeper understanding of cetacean evolution and facilitate informed conservation efforts for this enigmatic species.

## Data availability Statement

The base-called sequence data, assembly, and annotations have been deposited at the Aotearoa Genomic Database Repository (https://data.agdr.org.nz/). The data will be made available for scientific and conservation-related purposes on behalf of Ōraka-Aparima Rūnaka, with requests for commercial applications deferred to them. A github repository outlining the steps of the assembly and the code and data necessary for plotting the figures presented here is available (https://github.com/osilander/hourglass-assembly).

## Acknowledgments

This project was made possible with the assistance of kaimahi representing the Rūnaka of Ōraka-Aparima, including Riki Dallas and Iain MacCallum. We thank the Rūnaka for allowing us to access their taonga and for collaborating with us to codesign this kaupapa. We also thank the people who alerted us about the stranding, including Sonia Rahiti; Department of Conservation staff; staff and postgraduates of the Cetacean Ecology Research Group (CERG) and Massey University for their support with dissection and postmortem sampling. We further thank Melissa Nehmens for feedback on methods.

## Author Contributions

NM assisted in DNA isolation and performed DNA sequencing; JLR performed DNA isolation and assisted in DNA sequencing; AW assisted in genome assembly, analyses, and in drafting the manuscript; AA assisted in drafting manuscript and communication with Ōraka-Aparima; ROS supported the dissection and facilitated ongoing discussions with Ōraka-Aparima; MJ provided materials, input on data availability, and guidance from Ōraka-Aparima; KS performed dissection, tissue isolation, and manuscript drafting; OKS performed genome assembly, analyses, and drafted the manuscript; all authors edited and approved the manuscript.

## Supplementary Tables and Figures

**Supplementary Table 1.** List of other Delphinoidea genomes.

| Organism | Assembly ID | Date | Contigs | Technology |
| --- | --- | --- | --- | --- |
| *Tursiops aduncus* | GCA_003227395.1 | Jun 14, 2018 | 44,281 | Illumina HiSeq; Illumina NovaSeq |
| *Lagenorhynchus obliquidens* | GCA_003676395.1 | Oct 23, 2018 | 21,792 | Illumina HiSeq; Illumina NovaSeq |
| *Phocoena sinus* | GCA_008692025.1 | Sep 26, 2019 | 272 | PacBio Sequel I; Illumina NovaSeq; Arima Genomics Hi-C; Bionano Genomics DLE-1 |
| *Tursiops truncatus* | GCA_011762595.1 | Mar 27, 2020 | 1,035 | PacBio Sequel I CLR; Illumina NovaSeq; Arima Genomics Hi-C; Bionano Genomics DLS |
| *Orcinus orca* | GCA_937001465.1 | May 3, 2022 | 570 | PacBio and Hi-C |
| *Lagenorhynchus albirostris* | GCA_949774975.1 | Apr 8, 2023 | 1,479 | PacBio and Arima2 Hi-C |
| *Delphinus delphis* | GCA_949987515.1 | May 1, 2023 | 1,598 | PacBio and Arima2 Hi-C |
| *Stenella coeruleoalba* | GCA_951394435.1 | Jun 16, 2023 | 1,630 | PacBio and Arima2 Hi-C |
| *Globicephala melas* | GCA_963455315.1 | Sep 21, 2023 | 2,076 | PacBio and Arima2 Hi-C |

**Supplementary Table 2.** BUSCO completeness in Laurasiatheria across basecallers and assemblers.

| Assembler | Basecaller | Single copy | Duplicate | Fragmented | Missing |
| --- | --- | --- | --- | --- | --- |
| Goldrush | Guppy | 11,795 (96.4%) | 210 (1.72%) | 98 (0.80%) | 131 (1.07%) |
| Goldrush | Dorado | 11,826 (96.7%) | 191 (1.56%) | 92 (0.75%) | 124 (1.01%) |
| Nextdenovo | Guppy | 11,937 (97.6%) | 118 (0.96%) | 82 (0.67%) | 96 (0.78%) |
| Nextdenovo | Dorado | 11,936 (97.6%) | 116 (0.95%) | 78 (0.64%) | 104 (0.85%) |
| Raven | Guppy | 12,017 (98.2%) | 109 (0.89%) | 58 (0.47%) | 50 (0.41%) |
| Raven | Dorado | 12,024 (98.3%) | 107 (0.87%) | 55 (0.45%) | 48 (0.39%) |
| Raven | Dorado Q16 | 12,016 (98.2%) | 104 (0.85%) | 58 (0.47%) | 56 (0.46%) |

**Supplementary Table 3.** BUSCO completeness of the polished Dorado-basecalled Raven hourglass assembly relative to four mammalian lineages and to all eukaryotes.

| Lineage | Single copy and complete | Duplicated | Fragmented | Missing |
| --- | --- | --- | --- | --- |
| Laurasiatheria | 12,056 (98.55%) | 96 (0.78%) | 34 (0.28%) | 48 (0.39%) |
| Cetartiodactyla | 13,118 (98.37%) | 118 (0.88%) | 67 (0.50%) | 32 (0.24%) |
| Eutheria | 11,171 (98.29%) | 75 (0.66%) | 49 (0.43%) | 70 (0.62%) |
| Mammalia | 9,088 (98.50%) | 63 (0.68%) | 34 (0.37%) | 41 (0.45%) |
| Eukarya | 250 (98.04%) | 5 (1.96%) | 0 (0%) | 0 (0%) |

**Supplementary Figure 1.**
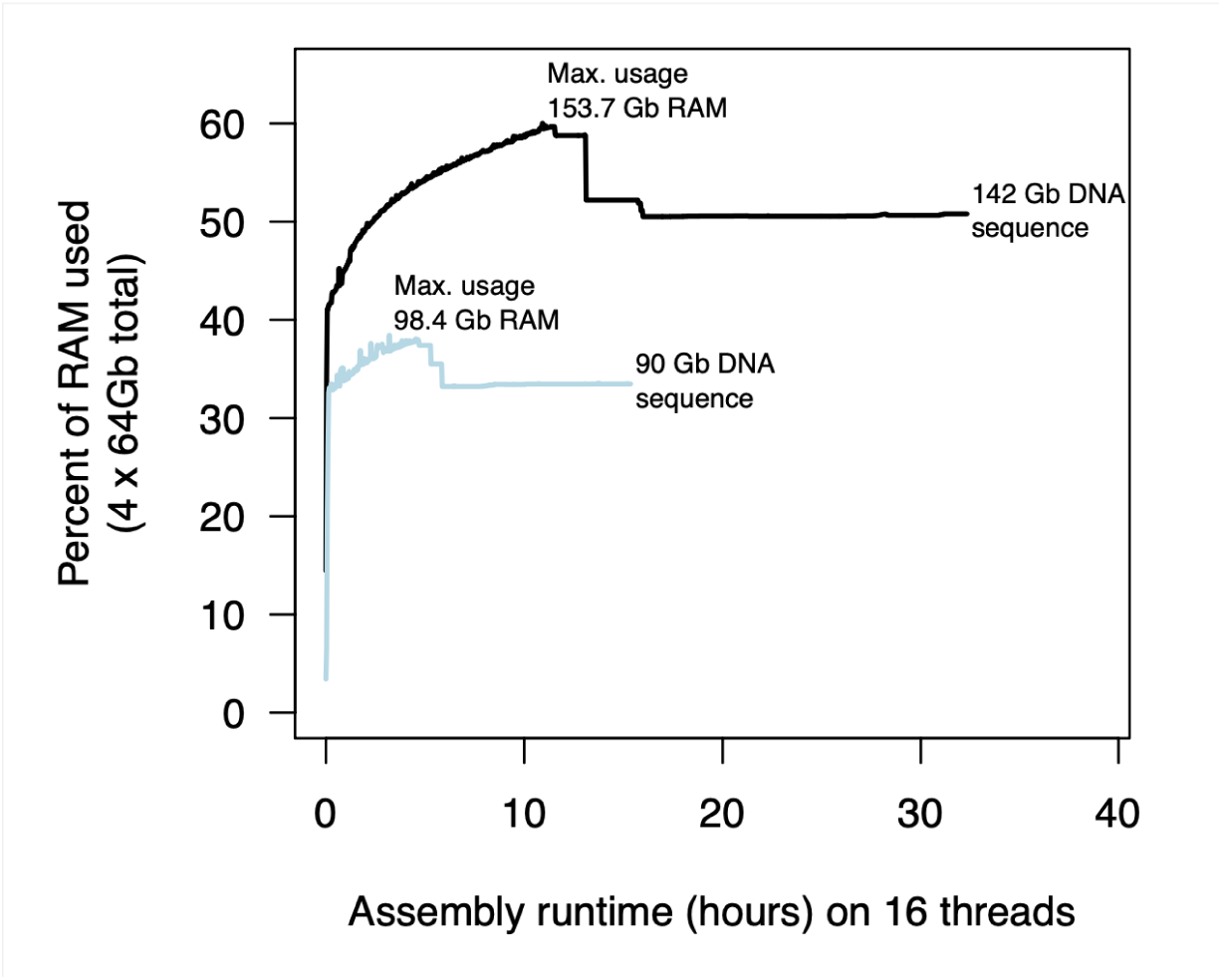
RAM usage during Raven genome assembly. The plot shows the amount of RAM used by Raven when performing assembly with 16 threads and using sequencing datasets with either 142 Gbp (the full dataset presented here) or 90 Gbp (a subset filtered by prioritising longer reads). The assembly from the 142 Gbp dataset required a maximum of 153.7 Gb of RAM, beyond the scale of most consumer laptops. The initial assembly from 142 Gbp data had 1,243 contigs, 2.44 Gbp total size, and an N50 of 8.08 Mbp. The initial assembly from the 90 Gbp dataset was slightly lower quality (1,621 contigs, 2.47 Gbp total length, N50 5.85 Mbp), but required less than 100 Gb total RAM. Current high end laptops have up to 128 Gb of RAM (for example, the HP ZBook Fury 16 G10, configurable with 128 Gb RAM and an A1000 GPU for less than $3,000 USD).

**Supplementary Figure 2.**
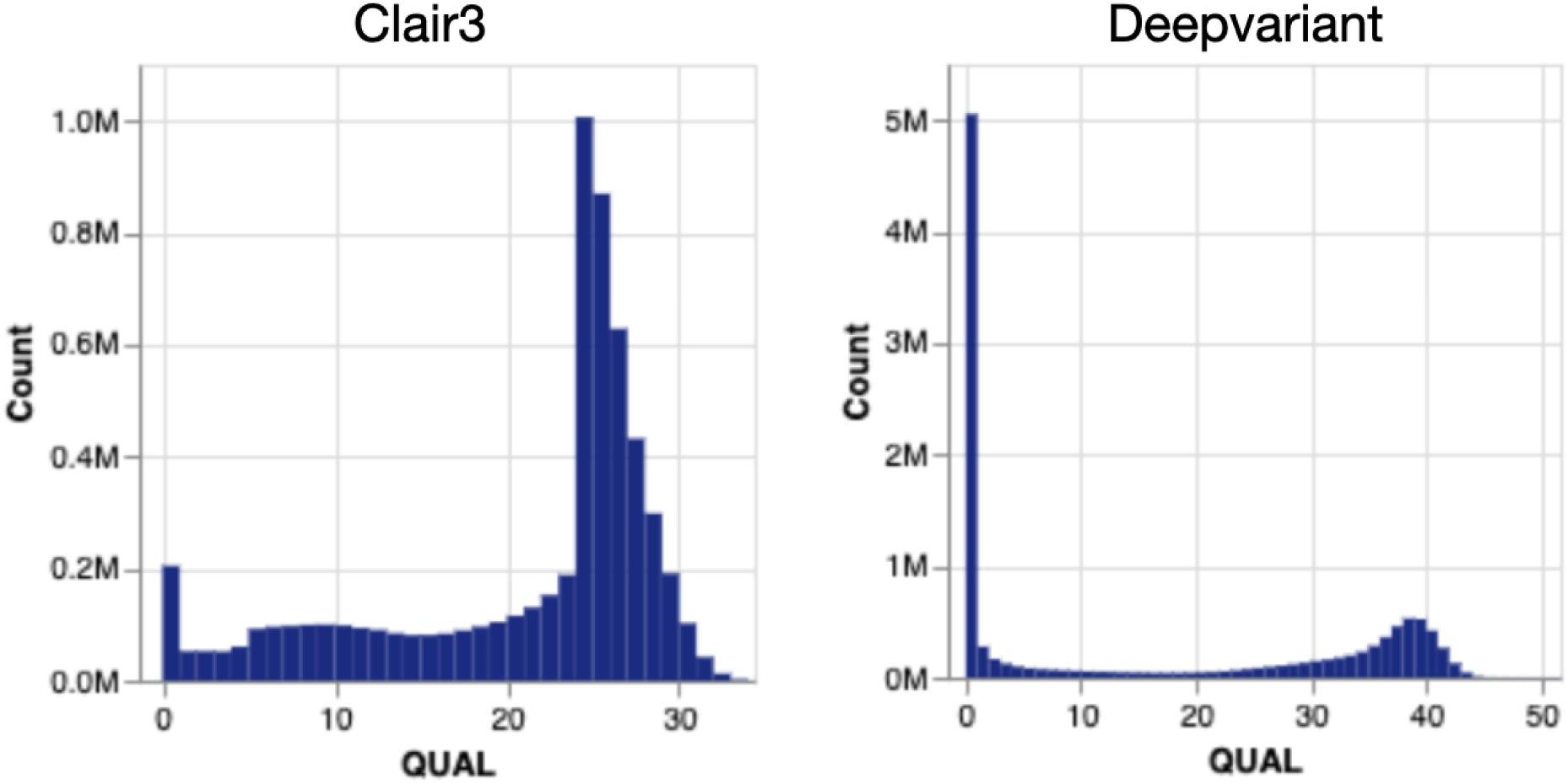
Genotype quality scores for Clair3 and DeepVariant. Clair3 calls have a minimum around q14; Deepvariant at around q20. We filtered both callsets to include only variants with higher qualities than this, respectively, and then merged the two filtered callsets to form a set of high quality genotype calls.

**Supplementary Figure 3.**
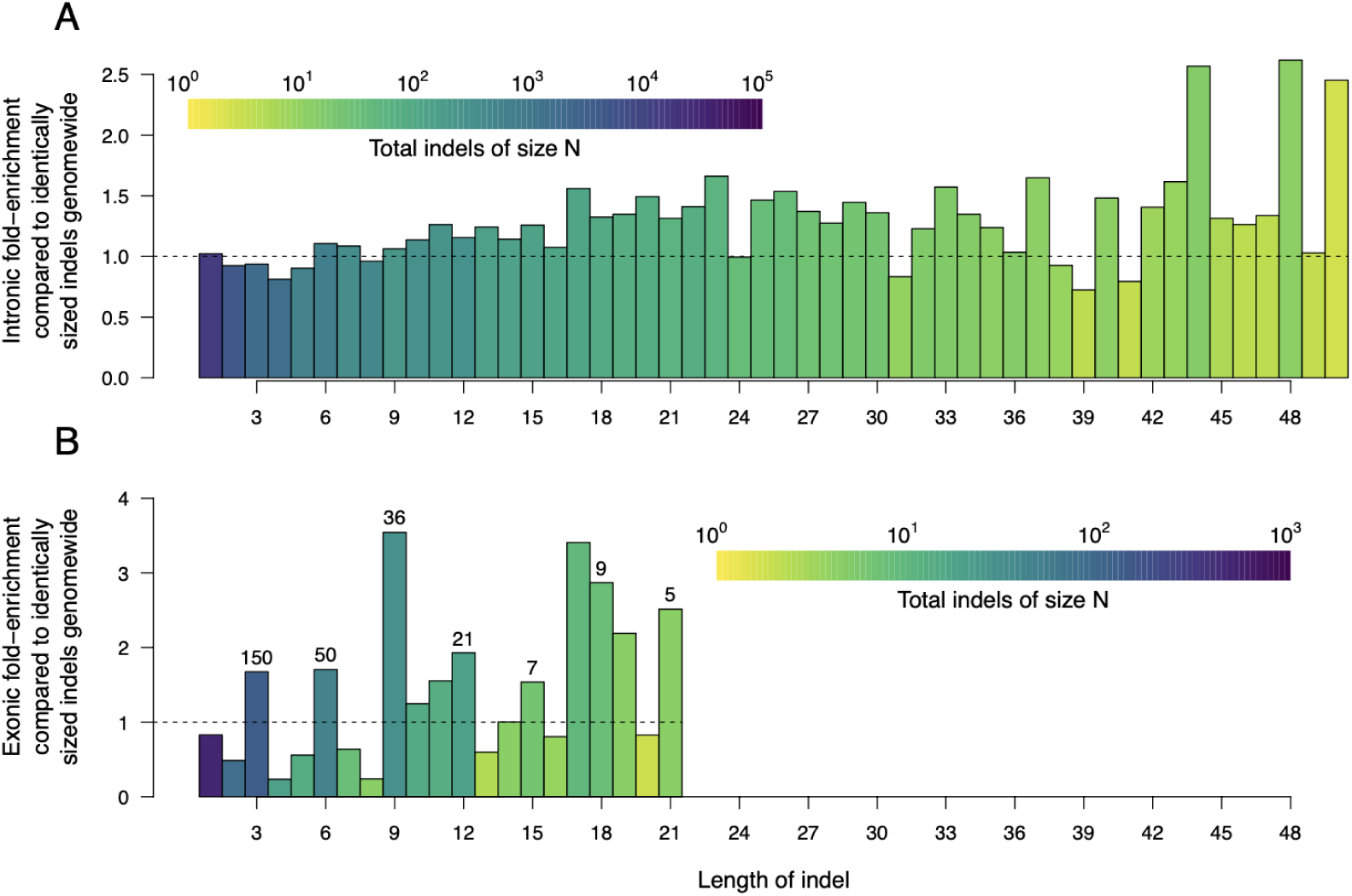
Enrichment and depletion of specific indel sizes in introns and exons relative to all genomic locations. A. The ratio of the number of indels of each size in introns relative to the number of indels of that size genome-wide. If there is no depletion or enrichment, the expected ratio is one (dotted line). Instead, there is a slight depletion of small intronic indels (two to five bp) and an enrichment of larger intronic indels (between five and 25 bp). Overall, we found a slight enrichment of indels in introns compared to the remainder of the genome (7.21% observed vs. 6.60% expected). **B**. The analogous plot as in (A) but for exonic regions. Here there is a specific depletion of indels that are not multiples of three bp. This is expected, as indels that are not multiples of three change the reading frame. For both panels, the colour of the bar indicates the total number of indels at that size (on a log scale). For panel (B), above each bar that is a multiple of three, we show the total number of indels that are of that size. For example, we observed a total of 150 three bp indels in exons. Relative to the total genome-wide number of three bp deletions and normalising for the fraction of indels in exonic regions, we would have expected approximately 1/2 that number (dotted line). The total fraction of indels across all sizes in exonic and intronic regions must sum to one; thus, if some indel sizes are depleted, others must necessarily be enriched. We do not show the relative depletion or enrichment for exonic indels beyond 21 bp as these are small in number and fold-enrichment or depletion varies widely. Note that both the y-axes scales and colour scales are different on each plot.

**Supplementary Figure 4.**
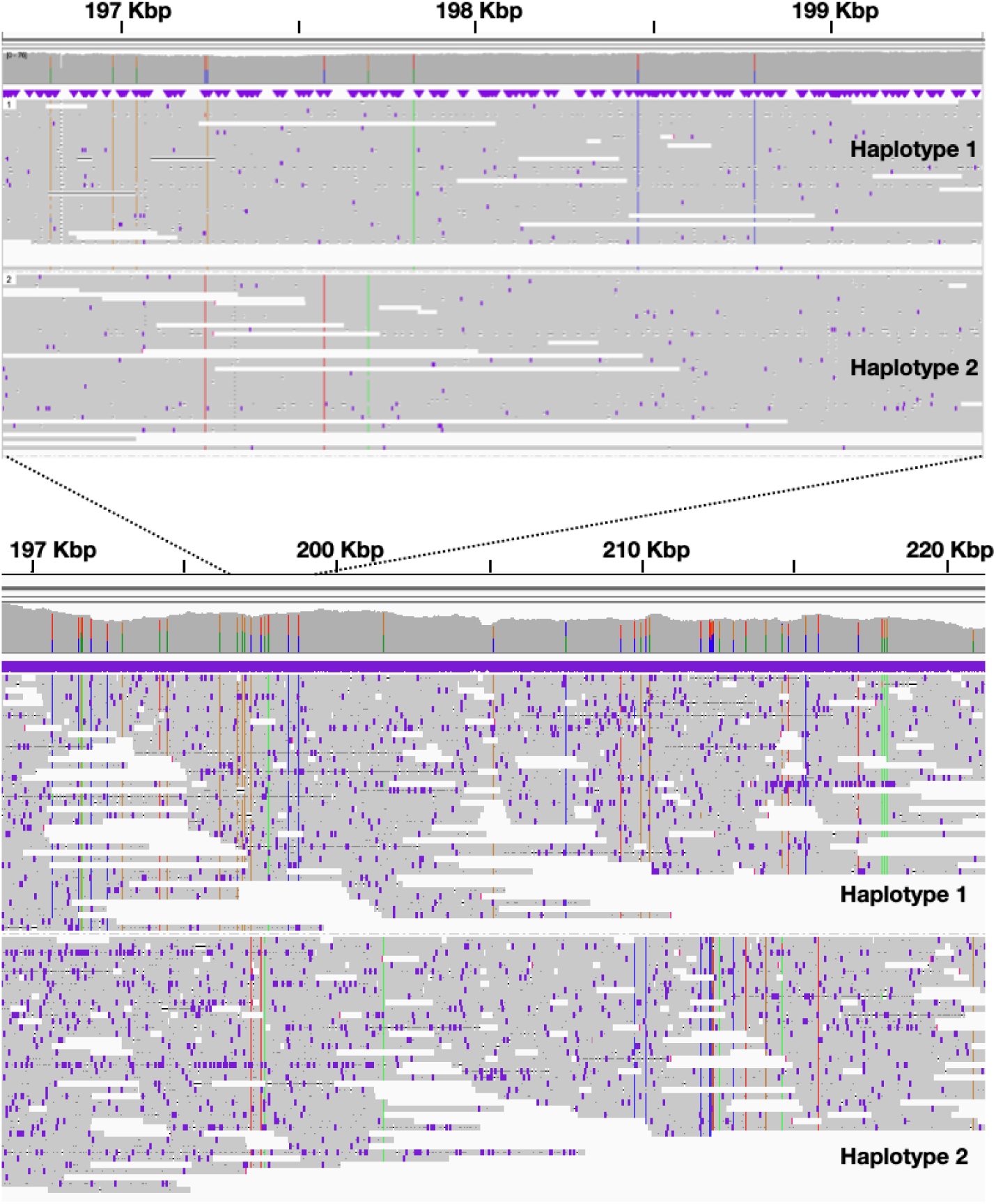
Phased haplotigs at 1 Kbp (top) and 10 Kbp (bottom) scales. Long Oxford Nanopore 10.4.1 chemistry reads allow phasing across the genome, despite relatively low levels of polymorphism in some regions (e.g. fewer than ten SNPs in the 10 kbp region between 200 Kbp and 210 Kbp above).

## References

Acevedo, Jorge, Sarah Garthe, and Alejandro González. 2017. “First Sighting of a Live Hourglass Dolphin (Lagenorhynchus Cruciger) in Inland Waters of Southern Chile.” Polar Biology 40 (2): 483–86.

Banguera-Hinestroza, E., P. G. H. Evans, L. Mirimin, R. J. Reid, B. Mikkelsen, A. S. Couperus, R. Deaville, E. Rogan, and A. R. Hoelzel. 2014. “Phylogeography and Population Dynamics of the White-Sided Dolphin (Lagenorhynchus Acutus) in the North Atlantic.” Conservation Genetics 15 (4): 789–802.

Bogdahn, Ingvar. 2015. “Agriculture-Independent, Sustainable, Fail-Safe and Efficient Food Production by Autotrophic Single-Cell Protein,” August. 10.7287/PEERJ.PREPRINTS.1279V2.

Brůna, Tomáš, Katharina J. Hoff, Alexandre Lomsadze, Mario Stanke, and Mark Borodovsky. 2021. “BRAKER2: Automatic Eukaryotic Genome Annotation with GeneMark-EP+ and AUGUSTUS Supported by a Protein Database.” NAR Genomics and Bioinformatics 3 (1): lqaa108.

Brůna, Tomáš, Alexandre Lomsadze, and Mark Borodovsky. 2020. “GeneMark-EP+: Eukaryotic Gene Prediction with Self-Training in the Space of Genes and Proteins.” NAR Genomics and Bioinformatics 2 (2): lqaa026.

Buchfink, Benjamin, Chao Xie, and Daniel H. Huson. 2015. “Fast and Sensitive Protein Alignment Using DIAMOND.” Nature Methods 12 (1): 59–60.

Cleary, John G., Ross Braithwaite, Kurt Gaastra, Brian S. Hilbush, Stuart Inglis, Sean A. Irvine, Alan Jackson, et al. 2014. “Joint Variant and de Novo Mutation Identification on Pedigrees from High-Throughput Sequencing Data.” Journal of Computational Biology: A Journal of Computational Molecular Cell Biology 21 (6): 405–19.

Cope, E. D. 1866. “Third Contribution to the History of the Balaenidae and Delphinidae.” Proceedings of the Academy of Natural Sciences of. https://www.jstor.org/stable/4059707.

Danecek, Petr, James K. Bonfield, Jennifer Liddle, John Marshall, Valeriu Ohan, Martin O. Pollard, Andrew Whitwham, et al. 2021. “Twelve Years of SAMtools and BCFtools.” GigaScience 10 (2). 10.1093/gigascience/giab008.

De Coster, Wouter, and Rosa Rademakers. 2023. “NanoPack2: Population-Scale Evaluation of Long-Read Sequencing Data.” Bioinformatics 39 (5). 10.1093/bioinformatics/btad311.

Dellabianca, Natalia, Gabriela Scioscia, Adrián Schiavini, and Andrea Raya Rey. 2012. “Occurrence of Hourglass Dolphin (Lagenorhynchus Cruciger) and Habitat Characteristics along the Patagonian Shelf and the Atlantic Ocean Sector of the Southern Ocean.” Polar Biology 35 (12): 1921–27.

Dorado. 2023. Github. https://github.com/nanoporetech/dorado.

Dutoit, Ludovic, Kieren J. Mitchell, Nicolas Dussex, Catherine M. Kemper, Petter Larsson, Love Dalén, Nicolas J. Rawlence, and Felix G. Marx. 2022. “Convergent Evolution of Skim Feeding in Baleen Whales.” bioRxiv. 10.1101/2022.08.16.504064.

Fernandez, Mercedes, Barbara Beron-Vera, Nestor A. Garcia, J. Antonio Raga, and Enrique A. Crespo. 2003. “Notes: FOOD AND PARASITES FROM TWO HOURGLASS DOLPHINS, LAGENORHYNCHUS CRUCIGER (QUOY AND GAIMARD, 1824), FROM PATAGONIAN WATERS.” Marine Mammal Science 19 (4): 832–36.

Garrison, Nanibaa’ A., Māui Hudson, Leah L. Ballantyne, Ibrahim Garba, Andrew Martinez, Maile Taualii, Laura Arbour, Nadine R. Caron, and Stephanie Carroll Rainie. 2019. “Genomic Research Through an Indigenous Lens: Understanding the Expectations.” Annual Review of Genomics and Human Genetics 20 (August): 495–517.

Goodall, R. N. P. 1997. “Review of Sightings of the Hourglass Dolphin, Lagenorhynchus Cruciger, in the South American Sector of the Antarctic and Sub-Antarctic.” Report of the International Whaling Commission.

Goodall, R. N. P., A. N. Baker, P. B. Best, M. Meyer, and N. Miyazaki. 1997. “On the Biology of the Hourglass Dolphin, Lagenorhynchus Cruciger (Quoy and Gaimard, 1824).” Annual Report of the International Whaling Commission 47: 985–99.

Gotoh, Osamu. 2008. “A Space-Efficient and Accurate Method for Mapping and Aligning cDNA Sequences onto Genomic Sequence.” Nucleic Acids Research 36 (8): 2630–38.

Gurevich, Alexey, Vladislav Saveliev, Nikolay Vyahhi, and Glenn Tesler. 2013. “QUAST: Quality Assessment Tool for Genome Assemblies.” Bioinformatics 29 (8): 1072–75.

Hall, Michael B., Ryan R. Wick, Louise M. Judd, An N. T. Nguyen, Eike J. Steinig, Ouli Xie, Mark R. Davies, Torsten Seemann, Timothy P. Stinear, and Lachlan J. M. Coin. 2024. “Benchmarking Reveals Superiority of Deep Learning Variant Callers on Bacterial Nanopore Sequence Data.” bioRxiv. 10.1101/2024.03.15.585313.

Harlin-Cognato, April D., and Rodney L. Honeycutt. 2006. “Multi-Locus Phylogeny of Dolphins in the Subfamily Lissodelphininae: Character Synergy Improves Phylogenetic Resolution.” BMC Evolutionary Biology 6 (November): 87.

Hassanin, Alexandre, Frédéric Delsuc, Anne Ropiquet, Catrin Hammer, Bettine Jansen van Vuuren, Conrad Matthee, Manuel Ruiz-Garcia, et al. 2012. “Pattern and Timing of Diversification of Cetartiodactyla (Mammalia, Laurasiatheria), as Revealed by a Comprehensive Analysis of Mitochondrial Genomes.” Comptes Rendus Biologies 335 (1): 32–50.

Hoff, Katharina J., Simone Lange, Alexandre Lomsadze, Mark Borodovsky, and Mario Stanke. 2016. “BRAKER1: Unsupervised RNA-Seq-Based Genome Annotation with GeneMark-ET and AUGUSTUS.” Bioinformatics 32 (5): 767–69.

Hoff, Katharina J., Alexandre Lomsadze, Mark Borodovsky, and Mario Stanke. 2019. “Whole-Genome Annotation with BRAKER.” Methods in Molecular Biology 1962: 65–95.

Huang, Neng, and Heng Li. 2023. “Compleasm: A Faster and More Accurate Reimplementation of BUSCO.” *Bioinformatics*, September. 10.1093/bioinformatics/btad595.

Hu, Jiang, Zhuo Wang, Zongyi Sun, Benxia Hu, Adeola Oluwakemi Ayoola, Fan Liang, Jingjing Li, et al. 2023. “An Efficient Error Correction and Accurate Assembly Tool for Noisy Long Reads.” bioRxiv. 10.1101/2023.03.09.531669.

Iwata, Hiroaki, and Osamu Gotoh. 2012. “Benchmarking Spliced Alignment Programs Including Spaln2, an Extended Version of Spaln That Incorporates Additional Species-Specific Features.” Nucleic Acids Research 40 (20): e161.

Jennings, Lydia, Talia Anderson, Andrew Martinez, Rogena Sterling, Dominique David Chavez, Ibrahim Garba, Maui Hudson, Nanibaa’ A. Garrison, and Stephanie Russo Carroll. 2023. “Applying the ‘CARE Principles for Indigenous Data Governance’ to Ecology and Biodiversity Research.” Nature Ecology & Evolution 7 (10): 1547–51.

Kuznetsov, Dmitry, Fredrik Tegenfeldt, Mosè Manni, Mathieu Seppey, Matthew Berkeley, Evgenia V. Kriventseva, and Evgeny M. Zdobnov. 2023. “OrthoDB v11: Annotation of Orthologs in the Widest Sampling of Organismal Diversity.” Nucleic Acids Research 51 (D1): D445–51.

Kyhn, Line A., J. Tougaard, F. Jensen, M. Wahlberg, G. Stone, A. Yoshinaga, K. Beedholm, and P. T. Madsen. 2009. “Feeding at a High Pitch: Source Parameters of Narrow Band, High-Frequency Clicks from Echolocating off-Shore Hourglass Dolphins and Coastal Hector’s Dolphins.” The Journal of the Acoustical Society of America 125 (3): 1783–91.

Leduc, R. G., W. F. Perrin, and A. E. Dizon. 1999. “Phylogenetic Relationships Among the Delphinid Cetaceans Based on Full Cytochrome B Sequences.” Marine Mammal Science 15 (3): 619–48.

Lee, Kyunglee, Junmo Lee, Hawsun Sohn, Yuna Cho, Young-Min Choi, Hye Kwon Kim, Ji Hyung Kim, and Dae Gwin Jeong. 2018. “Complete Mitochondrial Genome of the Pacific White-Sided Dolphin Lagenorhynchus Obliquidens (Cetacea: Delphinidae).” Conservation Genetics Resources 10 (2): 201–4.

Li, Heng. 2018. “Minimap2: Pairwise Alignment for Nucleotide Sequences.” Bioinformatics 34 (18): 3094–3100.

Lin, Yunzhi, Chen Ye, Xingzhu Li, Qinyao Chen, Ying Wu, Feng Zhang, Rui Pan, et al. 2023. “quarTeT: A Telomere-to-Telomere Toolkit for Gap-Free Genome Assembly and Centromeric Repeat Identification.” Horticulture Research 10 (8): uhad127.

Lomsadze, Alexandre, Vardges Ter-Hovhannisyan, Yury O. Chernoff, and Mark Borodovsky. 2005. “Gene Identification in Novel Eukaryotic Genomes by Self-Training Algorithm.” Nucleic Acids Research 33 (20): 6494–6506.

MacLeod, C. D. 2009. “Global Climate Change, Range Changes and Potential Implications for the Conservation of Marine Cetaceans: A Review and Synthesis.” Endangered Species Research 7 (June): 125–36.

Marçais, Guillaume, and Carl Kingsford. 2011. “A Fast, Lock-Free Approach for Efficient Parallel Counting of Occurrences of K-Mers.” Bioinformatics 27 (6): 764–70.

Marchesi, M., Lida E. Pimper, M. S. Mora, and R. Goodall. 2016. “The Vertebral Column of the Hourglass Dolphin (Lagenorhynchus Cruciger, Quoy and Gaimard, 1824), with Notes on Its Functional Properties in Relation to Its Habitat.” Aquatic Mammals 42 (September): 306–16.

Martin, Marcel, Murray Patterson, Shilpa Garg, Sarah O. Fischer, Nadia Pisanti, Gunnar W. Klau, Alexander Schöenhuth, and Tobias Marschall. 2016. “WhatsHap: Fast and Accurate Read-Based Phasing.” bioRxiv. 10.1101/085050.

Mc Cartney, Ann M., Jane Anderson, Libby Liggins, Maui L. Hudson, Matthew Z. Anderson, Ben TeAika, Janis Geary, Robert Cook-Deegan, Hardip R. Patel, and Adam M. Phillippy. 2022. “Balancing Openness with Indigenous Data Sovereignty: An Opportunity to Leave No One behind in the Journey to Sequence All of Life.” Proceedings of the National Academy of Sciences of the United States of America 119 (4). 10.1073/pnas.2115860119.

McGowen, Michael R. 2011. “Toward the Resolution of an Explosive Radiation--a Multilocus Phylogeny of Oceanic Dolphins (Delphinidae).” Molecular Phylogenetics and Evolution 60 (3): 345–57.

Medaka. 2023. Github. https://github.com/nanoporetech/medaka.

Morin, Phillip A., Alana Alexander, Mark Blaxter, Susana Caballero, Olivier Fedrigo, Michael C. Fontaine, Andrew D. Foote, et al. 2020. “Building Genomic Infrastructure: Sequencing Platinum-standard Reference-quality Genomes of All Cetacean Species.” Marine Mammal Science 36 (4): 1356–66.

Ondov, Brian D., Todd J. Treangen, Páll Melsted, Adam B. Mallonee, Nicholas H. Bergman, Sergey Koren, and Adam M. Phillippy. 2016. “Mash: Fast Genome and Metagenome Distance Estimation Using MinHash.” Genome Biology 17 (1): 132.

Peters, Katharina J., Sarah J. Bury, Bethany Hinton, Emma L. Betty, Déborah Casano-Bally, Guido J. Parra, and Karen A. Stockin. 2022. “Too Close for Comfort? Isotopic Niche Segregation in New Zealand’s Odontocetes.” Biology 11 (8). 10.3390/biology11081179.

Poplin, Ryan, Pi-Chuan Chang, David Alexander, Scott Schwartz, Thomas Colthurst, Alexander Ku, Dan Newburger, et al. 2018. “A Universal SNP and Small-Indel Variant Caller Using Deep Neural Networks.” Nature Biotechnology 36 (10): 983–87.

Quinlan, Aaron R., and Ira M. Hall. 2010. “BEDTools: A Flexible Suite of Utilities for Comparing Genomic Features.” Bioinformatics 26 (6): 841–42.

Roach, Michael J., Simon A. Schmidt, and Anthony R. Borneman. 2018. “Purge Haplotigs: Allelic Contig Reassignment for Third-Gen Diploid Genome Assemblies.” BMC Bioinformatics 19 (1): 460.

Robbins, Paul, Hilary Habeck Hunt, Francisco Pelegri, and Jonathan Gilbert. 2023. “Sovereign Genes: Wildlife Conservation, Genetic Preservation, and Indigenous Data Sovereignty.” Frontiers in Conservation Science 4. 10.3389/fcosc.2023.1099562.

Santora, Jarrod A. 2012. “Habitat Use of Hourglass Dolphins near the South Shetland Islands, Antarctica.” Polar Biology 35 (5): 801–6.

Shafin, Kishwar, Trevor Pesout, Pi-Chuan Chang, Maria Nattestad, Alexey Kolesnikov, Sidharth Goel, Gunjan Baid, et al. 2021. “Haplotype-Aware Variant Calling with PEPPER-Margin-DeepVariant Enables High Accuracy in Nanopore Long-Reads.” Nature Methods 18 (11): 1322–32.

Shen, Wei, Shuai Le, Yan Li, and Fuquan Hu. 2016. “SeqKit: A Cross-Platform and Ultrafast Toolkit for FASTA/Q File Manipulation.” PloS One 11 (10): e0163962.

Simão, Felipe A., Robert M. Waterhouse, Panagiotis Ioannidis, Evgenia V. Kriventseva, and Evgeny M. Zdobnov. 2015. “BUSCO: Assessing Genome Assembly and Annotation Completeness with Single-Copy Orthologs.” Bioinformatics 31 (19): 3210–12.

Stanke, Mario, Mark Diekhans, Robert Baertsch, and David Haussler. 2008. “Using Native and Syntenically Mapped cDNA Alignments to Improve de Novo Gene Finding.” Bioinformatics 24 (5): 637–44.

Stanke, Mario, Oliver Schöffmann, Burkhard Morgenstern, and Stephan Waack. 2006. “Gene Prediction in Eukaryotes with a Generalized Hidden Markov Model That Uses Hints from External Sources.” BMC Bioinformatics 7 (February): 62.

Te Aika, Ben, Libby Liggins, Claire Rye, E. Owen Perkins, Jun Huh, Rudiger Brauning, Tracey Godfery, and Michael A. Black. 2023. “Aotearoa Genomic Data Repository: An āhuru Mōwai for Taonga Species Sequencing Data.” *Molecular Ecology Resources*, September. 10.1111/1755-0998.13866.

Thiele, Deborah, Edwin T. Chester, and Peter C. Gill. 2000. “Cetacean Distribution off Eastern Antarctica (80--150 E) during the Austral Summer of 1995/1996.” Deep-Sea Research. Part II, Topical Studies in Oceanography 47 (12-13): 2543–72.

Thorvaldsdóttir, Helga, James T. Robinson, and Jill P. Mesirov. 2013. “Integrative Genomics Viewer (IGV): High-Performance Genomics Data Visualization and Exploration.” Briefings in Bioinformatics 14 (2): 178–92.

Tillich, Michael, Pascal Lehwark, Tommaso Pellizzer, Elena S. Ulbricht-Jones, Axel Fischer, Ralph Bock, and Stephan Greiner. 2017. “GeSeq - Versatile and Accurate Annotation of Organelle Genomes.” Nucleic Acids Research 45 (W1): W6–11.

Todd, Victoria L. G., and Laura D. Williamson. 2022. “Cetacean Distribution in Relation to Oceanographic Features at the Kerguelen Plateau.” Polar Biology 45 (1): 113–26.

Tougaard, J., and L. Kyhn. 2009. “Echolocation Sounds of Hourglass Dolphins (Lagenorhynchus Cruciger) Are Similar to the Narrow Band High-Frequency Echolocation Sounds of the Dolphin Genus Cephalorhynchus.” Marine Mammal Science 26 (May): 239–45.

Vaser, Robert, and Mile Šikić. 2021. “Time- and Memory-Efficient Genome Assembly with Raven.” Nature Computational Science 1 (5): 332–36.

Vollmer, Nicole L., Erin Ashe, Robert L. Brownell Jr, Frank Cipriano, James G. Mead, Randall R. Reeves, Melissa S. Soldevilla, and Rob Williams. 2019. “Taxonomic Revision of the Dolphin Genus Lagenorhynchus.” Marine Mammal Science 35 (3): 957–1057.

Wick, Ryan R., and P. Menzel. 2017. “Filtlong.” Available Online: Github. com/rrwick/Filtlong (accessed on 15 August 2021).

Wong, Johnathan, Lauren Coombe, Vladimir Nikolić, Emily Zhang, Ka Ming Nip, Puneet Sidhu, René L. Warren, and Inanç Birol. 2023. “Linear Time Complexity de Novo Long Read Genome Assembly with GoldRush.” Nature Communications 14 (1): 2906.

Zheng, Zhenxian, Shumin Li, Junhao Su, Amy Wing-Sze Leung, Tak-Wah Lam, and Ruibang Luo. 2022. “Symphonizing Pileup and Full-Alignment for Deep Learning-Based Long-Read Variant Calling.” Nature Computational Science 2 (12): 797–803.

Zhong, Z., L. Shiue, S. Kaplan, and T. de Lange. 1992. “A Mammalian Factor That Binds Telomeric TTAGGG Repeats in Vitro.” Molecular and Cellular Biology 12 (11): 4834–43.

